# The global distribution of paired eddy covariance towers

**DOI:** 10.1101/2023.03.03.530958

**Authors:** Paul C. Stoy, Housen Chu, Emma Dahl, Daniela S. Cala, Victoria Shveytser, Susanne Wiesner, Ankur R. Desai, Kimberly A. Novick

## Abstract

The eddy covariance technique has revolutionized our understanding of ecosystem-atmosphere interactions. Eddy covariance studies often use a “paired” tower design in which observations from nearby towers are used to understand how different vegetation, soils, hydrology, or experimental treatment shape ecosystem function and surface-atmosphere exchange. Paired towers have never been formally defined and their global distribution has not been quantified. We compiled eddy covariance tower information to find towers that could be considered paired. Of 1233 global eddy covariance towers, 692 (56%) were identified as paired by our criteria. Paired towers had cooler mean annual temperature (mean = 9.9 °C) than the entire eddy covariance network (10.5 °C) but warmer than the terrestrial surface (8.9 °C) from WorldClim 2.1, on average. The paired and entire tower networks had greater average soil nitrogen (0.57-0.58 g/kg) and more silt (36.0-36.4%) than terrestrial ecosystems (0.38 g/kg and 30.5%), suggesting that eddy covariance towers sample richer soils than the terrestrial surface as a whole. Paired towers existed in a climatic space that was more different from the global climate distribution sampled by the entire eddy covariance network, as revealed by an analysis of the Kullback-Leibler divergence, but the edaphic space sampled by the entire network and paired towers was similar. The lack of paired towers with available data across much of Africa, northern, central, southern, and western Asia, and Latin America with few towers in savannas, shrublands, and evergreen broadleaf forests point to key regions, ecosystems, and ecosystem transitions in need of additional research. Few if any paired towers study the flux of ozone and other atmospherically active trace gases at the present. By studying what paired towers measure – and what they do not – we can make infrastructural investments to further enhance the value of FLUXNET as it moves toward its fourth decade.

## Introduction

The eddy covariance technique has been indispensable for understanding the role that ecosystems play in the earth system by controlling the fluxes of water, carbon dioxide and other greenhouse gases, ozone and other atmospherically active gases, heat, and momentum with the atmosphere (Baldocchi, 2008; Clifton et al., 2020). As eddy covariance networks have grown, we have had more opportunities to understand how different ecosystems impact these fluxes to help us quantify the critical services that they provide (Novick et al., 2022). These studies often arise via a “paired” study design where observations from two or more nearby eddy covariance towers are compared to understand how different ecosystems behave when subject to similar climatic conditions (Baldocchi, 2014). To our understanding, paired towers have never been defined, and therefore their unique role in flux science has not been fully articulated.

### What are paired towers?

There is no formal definition of paired towers, and different applications may arrive at different conclusions. At a minimum we suggest that paired towers include at least two eddy covariance towers in an area with similar macro-climatic conditions but with different vegetation, soil types, hydrologic regimes, or experimental treatments. In this way the role of vegetation, soils, or hydrologic context (e.g., uplands, wetlands, or water bodies) can be separated from macro-climatic conditions in determining surface-atmosphere exchange. It is important to note that microclimate can vary substantially across ecosystem transitions (Baker et al., 2014), but these differences are a function of the ecosystems themselves such that one should expect microclimate to differ across paired ecosystems.

Existing paired-tower studies adopt different approaches for categorizing similarity in macro-climatic conditions. For example, some paired tower experiments used sites that were immediately adjacent (Bavin et al., 2009), separated by the extent of an agricultural field (Vick et al., 2016), or co-located to within < 0.5 km (Moore et al., 2021). Other studies have adopted more liberal definitions. For example, to investigate the impacts of reforestation on surface temperature, Zhang et al., (2020) relied on paired grassland and forest towers co-located to within 30 km, a scale at which one may assume that macro-climate does not vary appreciably under many circumstances and with a spatial extent similar to, for example, the ∼32 km North American Regional Reanalysis (NARR, Mesinger et al., 2006). Ultimately, what constitutes “similar” will depend on the goals and questions of a given study. For site tower pairs at distances (and/or elevations) where climate can be expected to vary, statistical analyses of differences between key climate drivers like radiation, temperature, precipitation, and/or vapor pressure deficit is probably necessary.

The application of the paired tower approach depends on the question at hand. For example, a study of the role of ecosystem transitions on near-surface turbulence requires towers in adjacent ecosystems (e.g. Detto et al., 2008), but studies of the impact of land management on surface fluxes at a regional scale could consider tower pairs to be further apart (e.g. Runkle et al., 2017). We also note that edaphic characteristics often differ quite dramatically across small spatial extents including within the flux footprint (Herbst et al., 2021; Levy et al., 2020; Rey-Sanchez et al., 2022; Tuovinen et al., 2019; Vanderborght et al., 2010). We discuss the importance of quantifying soil heterogeneity when interpreting tower observations by detailing examples of edaphic heterogeneity among paired towers in the analysis below.

### What can paired towers do?

Paired towers have been fundamental for understanding how land use and land management impact land surface fluxes and microclimate (Baldocchi, 2020). Some of the first applications of long-term eddy covariance measurements, like the BOREAS campaign (Margolis and Ryan, 1997), were made with comparative ecosystem studies in mind. Our understanding of the carbon and water cycle consequences of land cover change and land management have been shaped by the paired tower approach to quantify impacts of forest management (Desai et al., 2005; Goulden et al., 2006; Starr et al., 2016), ecological restoration (Hemes et al., 2019), ecological disturbances (Amiro, 2001; Kowalski et al., 2004; Rebane et al., 2019), agricultural management (Baker and Griffis, 2005; Chi et al., 2016; French et al., 2020; Moore et al., 2020; Runkle et al., 2018), and more. Our understanding of how ecosystem carbon and water cycles interact to determine the ecosystem water use efficiency has therefore also been shaped by paired tower studies (Anapalli et al., 2019; Stoy et al., 2008; Volik et al., 2021) and observations over multiple years have been used to quantify interactions between the carbon and water cycles across multiple scales in time (Novick et al., 2015).

Paired towers also help us elucidate how vegetation controls biophysical attributes of ecosystems like land surface temperature (Luyssaert et al., 2014; Zhang et al., 2020) and air temperature (Baldocchi and Ma, 2013; Novick and Katul, 2020), land surface albedo (Duman et al., 2021; J. Juang et al., 2007a), surface roughness (Burakowski et al., 2018), and soil heat flux (Sun et al., 2019). They help us understand how ecosystem structural attributes and transitions – including terrestrial-aquatic transitions – impact near-surface turbulence (Detto et al., 2008; Eder et al., 2013; Rey-Sánchez et al., 2017; Turner et al., 2019). Paired towers can also help us understand how changes in energy partitioning at the land surface impact atmospheric boundary layer dynamics (Eder et al., 2015; Paleri et al., 2022; Ueyama et al., 2020), including those related to precipitation processes (J. Juang et al., 2007b; J.-Y. Juang et al., 2007; Manoli et al., 2016; Vick et al., 2016).

Paired towers are often used to extrapolate surface fluxes to global and regional scales using data-driven approaches (Jung et al., 2009; Poe et al., 2020) and land surface models (Yuan et al., 2021), and can help identify the mechanisms necessary to improve such models (Vanden Broucke et al., 2015). The proximity of paired towers can challenge remote sensing and modeling efforts if they lie within the same pixel (Heinsch et al., 2006) because land cover change impacts surface fluxes across small spatial scales (Rohatyn et al., 2022). Ongoing efforts to quantify tower spatial representativeness are critical for scaling flux tower observations to larger scales in space (Chu et al., 2021; Volk et al., 2023). While paired towers at their simplest are limited to a small sample size compared to a typical replicate study, the high time frequency of EC sampling made over long continuous periods of times makes it possible to tease out differences in fluxes in ways that infrequent plot sampling across replicates cannot.

Paired towers are increasingly used to compare how different ecosystems regulate non-CO_2_ greenhouse gas fluxes including methane (Delwiche et al., 2021; Krauss et al., 2016; Matthes et al., 2014) and nitrous oxide (Voglmeier et al., 2020), but there have been fewer studies of atmospherically active trace gases like ozone and NO_x_ from a paired flux tower perspective, which leaves critical gaps in our understanding. Paired tower observations also add value to eddy covariance measurements themselves by providing an avenue to quantify their uncertainty (Hollinger and Richardson, 2005; Kessomkiat et al., 2013; Lasslop et al., 2008; Oren et al., 2006; Post et al., 2015; Richardson et al., 2012; Richardson and Hollinger, 2005) including the ways in which different ecosystems impact surface-atmosphere energy balance closure (Anderson and Wang, 2014; Barr et al., 2006; Stoy et al., 2006). A major motivation of the present analysis is to encourage researchers to apply the paired tower approach to ask novel research questions going forward.

### Where do researchers have more (or fewer) opportunities to study paired towers?

Ecosystems across some parts of the globe simply have more eddy covariance towers than others (Chu et al., 2017; Pastorello et al., 2020; Xiao et al., 2012), leaving large swaths of the planet understudied. As a subset of the FLUXNET network, one can assume that paired towers share this geographic bias. We suspect that paired towers have an even more biased geographic distribution than the entire global eddy covariance network itself as they are likely to be in areas that are already well-studied, but any differences have not been quantified to date. Our goal in the present analysis is to quantify the geographic regions, climatic zones, and edaphic settings that have relatively more and fewer paired towers to characterize the current state of paired towers across the FLUXNET network and to identify areas in need of more research investment. We start by systematically identifying paired towers and then evaluate their spatial coverage and climatic and edaphic representativeness before discussing paired tower research going forward.

## Methods

### Identifying paired towers

We compiled locations of eddy covariance towers, climatology, and ecosystem type from the AmeriFlux, European Fluxes Database Cluster, AsiaFlux, OzFlux, and FLUXNET databases supplemented by information from synthesis manuscripts (e.g., Beringer et al., 2016; Cleverly et al., 2020). Data were accessed between September 2022 and January 2023. Towers were identified as paired by communication with principal investigators coupled with our interpretation of tower proximity and intent (e.g., if towers have a similar naming convention), tower observational periods, site descriptions, and site-relevant publications. We use a ∼32 km threshold as above to approximately delineate paired towers, keeping sites with pronounced elevational differences in mind. In this way we attempted to identify sites that have similar climatic conditions, on average. Consequently, the paired tower list that we generated (Appendix A) should be considered representative of eddy covariance towers that were likely designed with comparative ecosystem studies in mind, noting that the applicability of the paired tower approach depends on the question at hand as discussed above. We note that tower locations are continually changing as eddy covariance towers are built and decommissioned, and the results of the present analyses are subject to the ever-changing nature of the global eddy covariance network, listed in Appendix B. It is important to note that some paired towers are not included in flux networks for multiple reasons (e.g., Deshmukh et al., 2020). We exclude these sites from the present analysis, and we note a number of detailed studies of site representativeness across regional networks that provide key context for our global analysis (Hargrove et al., 2003; Sulkava et al., 2011; Villarreal et al., 2018; Villarreal and Vargas, 2021). An example of an area with multiple eddy covariance towers, all of which were installed with paired ecosystem studies in mind, is presented in Figure 1. From this perspective, the use of ‘paired’ relates more to the pairwise comparisons that can be made rather than towers themselves being a pair (i.e., implying two towers), and multiple intercomparisons can be made in intensively-instrumented regions with multiple towers.

**Figure 1:**
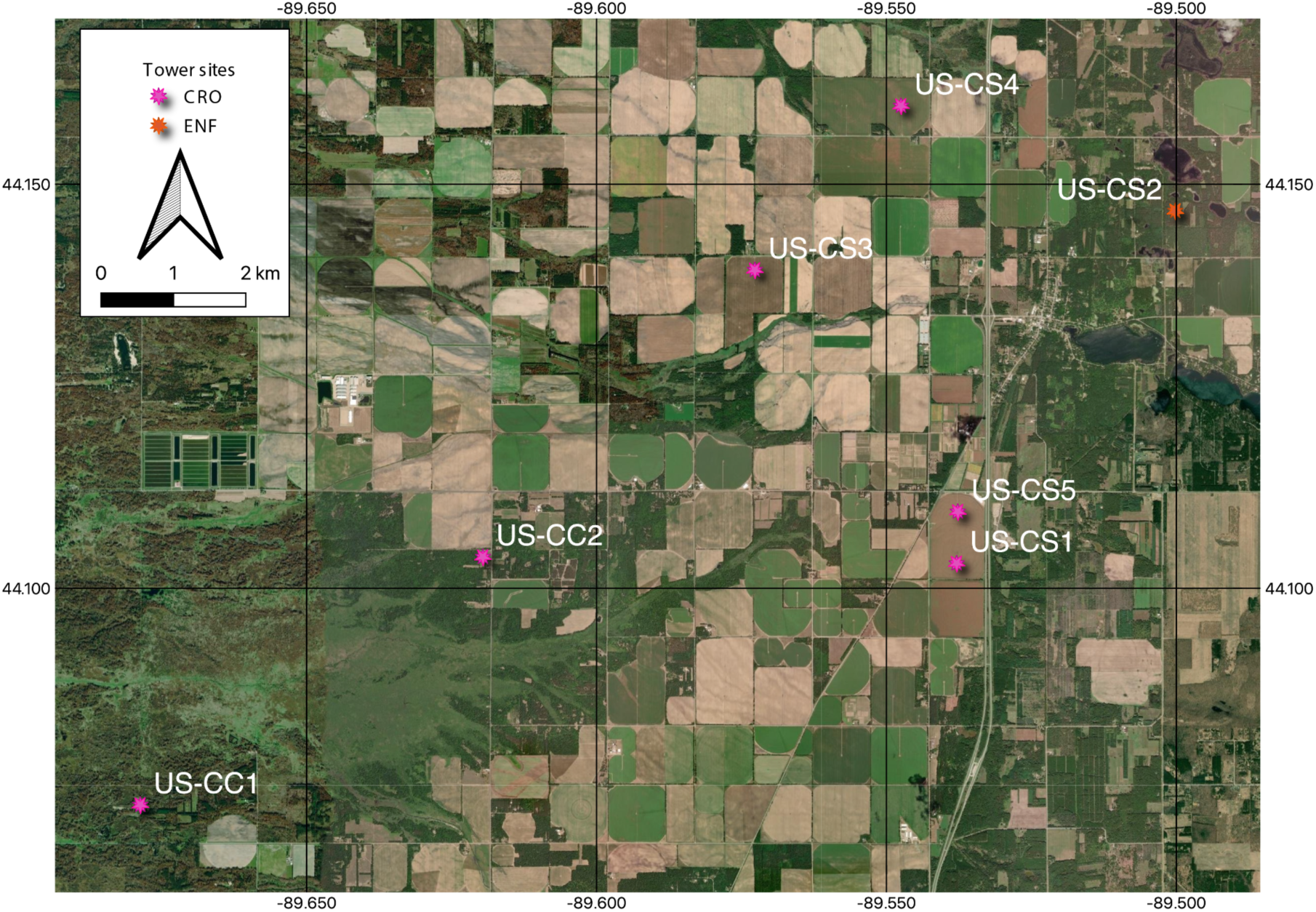
An example of a region with multiple paired towers in central Wisconsin, USA. Note that some towers (US-CS1 & US-CS5) exist within what could be considered a single field but were operational at different times (US-CS1: 2018-2019; US-CS5: 2021-onward) such that they would not be able to be directly compared under the same climatic conditions. Other towers were operational for short campaigns (e.g. US-CS4: 2020-2021; US-CC1 & US-CC2: 2021). The selection of tower pairs depends on the research question at hand. The base map is from QuickMapServices of QGIS.

### Global climate data

We seek to quantify the representativeness of eddy covariance tower placement with respect to global climatic and edaphic conditions. Mean annual temperature (MAT) and precipitation (MAP) data were available from the metadata of many tower sites, but not others. To approximate climatic conditions at these sites, and to add mean annual incident solar radiation (SRAD) to our analysis, we used observations from the WorldClim 2.1 database at ∼ 1 km resolution (Fick and Hijmans, 2017). WorldClim 2.1 data at tower locations were compiled using R (R Core Team, 2022) (Appendices A and B). Elevation data were likewise available for many towers, and when not was estimated using data from the *elevatr* package in R (Hollister et al., 2022) that accesses the Amazon Web Service Terrain Tiles, Open Topography Global Datasets, and USGS Elevation Point Query Service. We used a random draw of 10,000 data points on the sphere using the ‘runifsphere’ command in the *globe* package in R (Baddeley et al., 2017) to estimate the global distribution of terrestrial climatic conditions from WorldClim2.1 and elevation from *elevatr* for statistical analyses. We use tower-reported values for statistical analyses when possible and observations from global databases when necessary Information regarding each is available in the Appendices. Uncertainties in applying global database information instead of tower observations (Vuichard and Papale, 2015) are quantified below.

### Soil characteristics

Edaphic conditions are infrequently used to interpret eddy covariance observations despite the central role of soils in ecosystem functioning. We approximated edaphic conditions at each site using the SoilGrids 2.0 database (Poggio et al., 2021) assisted by the *soilDB* package (Beaudette et al., 2022) in R. We used data from the uppermost soil layer (0 - 5 cm) for simplicity and explored the mean estimated value of sand, silt, and clay fraction (in percent), pH, N concentration, soil organic matter content (in the fine soil fraction), bulk density, and cation exchange capacity. Observations from tower locations are compared against a global distribution of all quantities approximated by a random sample on the sphere (Baddeley et al., 2017) from 10,000 locations from SoilGrids 2.0. Distributions of global and eddy covariance tower sand, silt, and clay fractions were plotted in the soil texture triangle using the *soiltexture* package in R (Moeys, 2018).

### Statistical analysis: Kullback-Leibler divergence

We seek to quantify differences in the global distribution of climatic and edaphic variables versus the distribution sampled by all eddy covariance towers and the paired tower subset. To do so we use the Kullback-Liebler divergence (*D_KL_*, Kullback and Leibler, 1951) which measures the difference in information content between a reference distribution (here the random samples of global climate and soils data, *P*(*x*)) and a test distribution (here the climate and edaphic characteristics from the combined or paired eddy covariance sites, *Q*(*x*)):

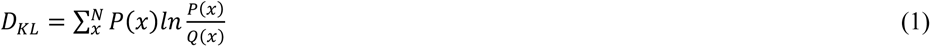

*D_KL_* values further from 0 indicate that the data distributions are more different. The *D_KL_* is sensitive to the number of bins (*N*) used in its calculation; we use *N* = 20 bins as a reasonable value that captures the distribution of the different datasets (Afgani et al., 2008). The distributions for some variables may be approximated using a statistical distribution (e.g., soil texture can be approximated by Beta distributions (Haskett et al., 1995) for which *D_KL_* can be calculated using their corresponding parameters (e.g. Stoy et al., 2022)), but we use the nonparametric *D_KL_* (equation 1) across all variables to aid in comparisons and to avoid parametric assumptions that may not be warranted.

### Statistical analysis: Wilcoxon rank sum tests

When testing for statistical differences between means it is critical to note that the paired tower and entire tower datasets are not independent; the paired tower network is a subset of all towers. We therefore compared paired towers and the *unpaired* tower network against the random draw of climate and elevation datasets as pairwise comparisons using the Wilcoxon rank sum test with continuity correction and applied the Holm-Bonferroni correction when interpreting statistical significance at the α < 0.01 level. Strictly speaking, the tower locations are a subset of the global climate and elevation datasets, but with a global terrestrial area of 148,326,000 km^2^ and data drawn from tower point locations or 1 km^2^ (or smaller) pixels, the chance of resampling is less than 1 in 100,000. We therefore assume that the tower data distributions and global data distributions are statistically distinct for the purposes of our study.

### Uncertainty

There are multiple sources of uncertainty when attempting to quantify where paired towers exist across geographic, climate, and edaphic space. These include uncertainties in identifying paired towers as noted, errors associated with compiling the ever-changing suite of tower locations (Appendices A and B), errors in soil and climate estimates from global databases including subpixel variability, uncertainty due to the extrapolation approach (Fick and Hijmans, 2017; Poggio et al., 2021), and uncertainties in tower measurements themselves including climatic and edaphic variability within the flux footprint and the representativeness of the flux footprint within the pixel dimensions (Chu et al., 2021). To help quantify the degree of uncertainty induced by the latter factors, we compare climate data reported from tower sites against that from WorldClim 2.1, noting that climate normals themselves of course are changing due to global climate change. We also study soil silt, sand, and clay content from the CHEESEHEAD19 database of 19 eddy covariance towers within a 10 × 10 km domain in northern Wisconsin, USA (Butterworth et al., 2021) using the average of three measurements from soil cores within the flux footprints (Hu et al., 2020), data from the NRCS soil survey (Shveytser et al., 2022), and data from SoilGrids 2.0 (Poggio et al., 2021) to demonstrate the range of soil texture values that may be encountered when using different datasets.

## Results

Of the 1233 eddy covariance towers that we compiled from global databases (Appendix B), we determined that 692 (56%) could be considered ‘paired’ (Appendix A). At the present, paired towers with publicly available data are absent from large expanses of the terrestrial biosphere including much of Africa, the Middle East, Central and South Asia, and Siberia (Figure 2). Paired towers in extremely cold ecosystems, and ecosystems with high average temperatures and high or low precipitation tend to be underrepresented in global databases (Figures 3-4), which is dominated by temperate ecosystems as also determined from the previous FLUXNET La Thuile synthesis dataset (Stoy et al., 2009; Xiao et al., 2012) and the FLUXNET2015 dataset (Pastorello et al., 2020). There are relatively few towers and fewer still paired towers in savannas, shrublands, evergreen broadleaf forests, snow-covered, and barren ecosystems than the global distribution of these IGBP categories (Table 1). Of regions with a higher density of eddy covariance measurements, North America has more towers (575) and paired towers (403, 70%) than Europe (396 and 190, 48%) but lower density of total towers (2.37 × 10^-5^ km^-2^ vs. 3.95 × 10^-5^ km^-2^) and paired towers (1.66 × 10^-5^ km^-2^ vs. 1.92 × 10^-5^ km^-2^, Table 2). The question remains: are paired towers located in less representative locations in climate and edaphic space than the eddy covariance network as a whole?

**Figure 2:**
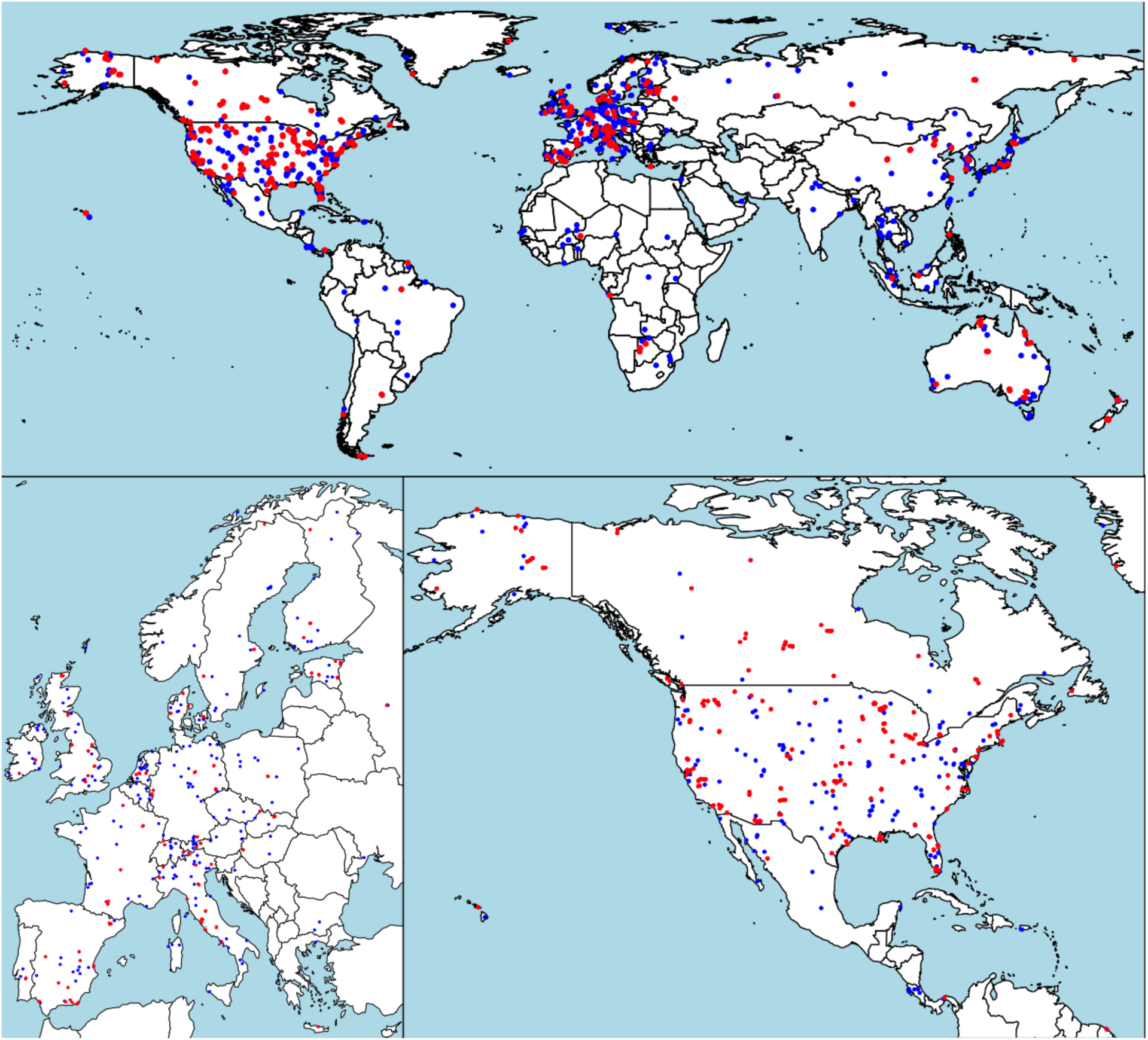
A map of the location of towers determined to be “paired” (red, Appendix A) with unpaired towers in blue (Appendix B) across the world excluding Antarctica (top), part of Europe (lower left), and part of the northern Americas (lower right).

**Figure 3:**
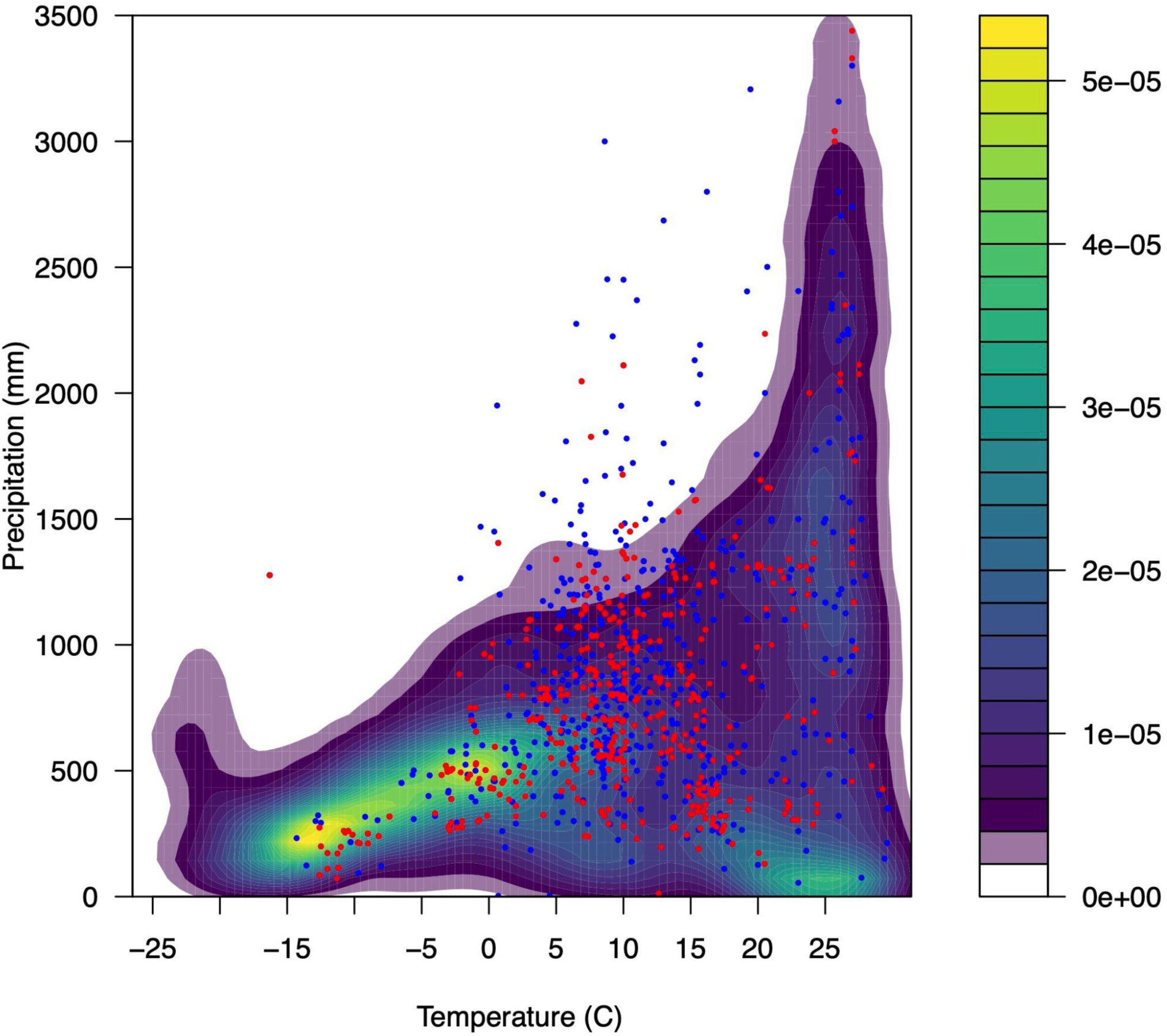
The mean annual temperature and precipitation of eddy covariance tower sites (blue dots) and paired towers (red dots) versus a two-dimensional kernel density estimate of global temperature and precipitation from the WorldClim 2.1 database (Fick and Hijmans, 2017). The probability of each pixel in global precipitation/temperature space is shown in the colorbar.

**Figure 4:**
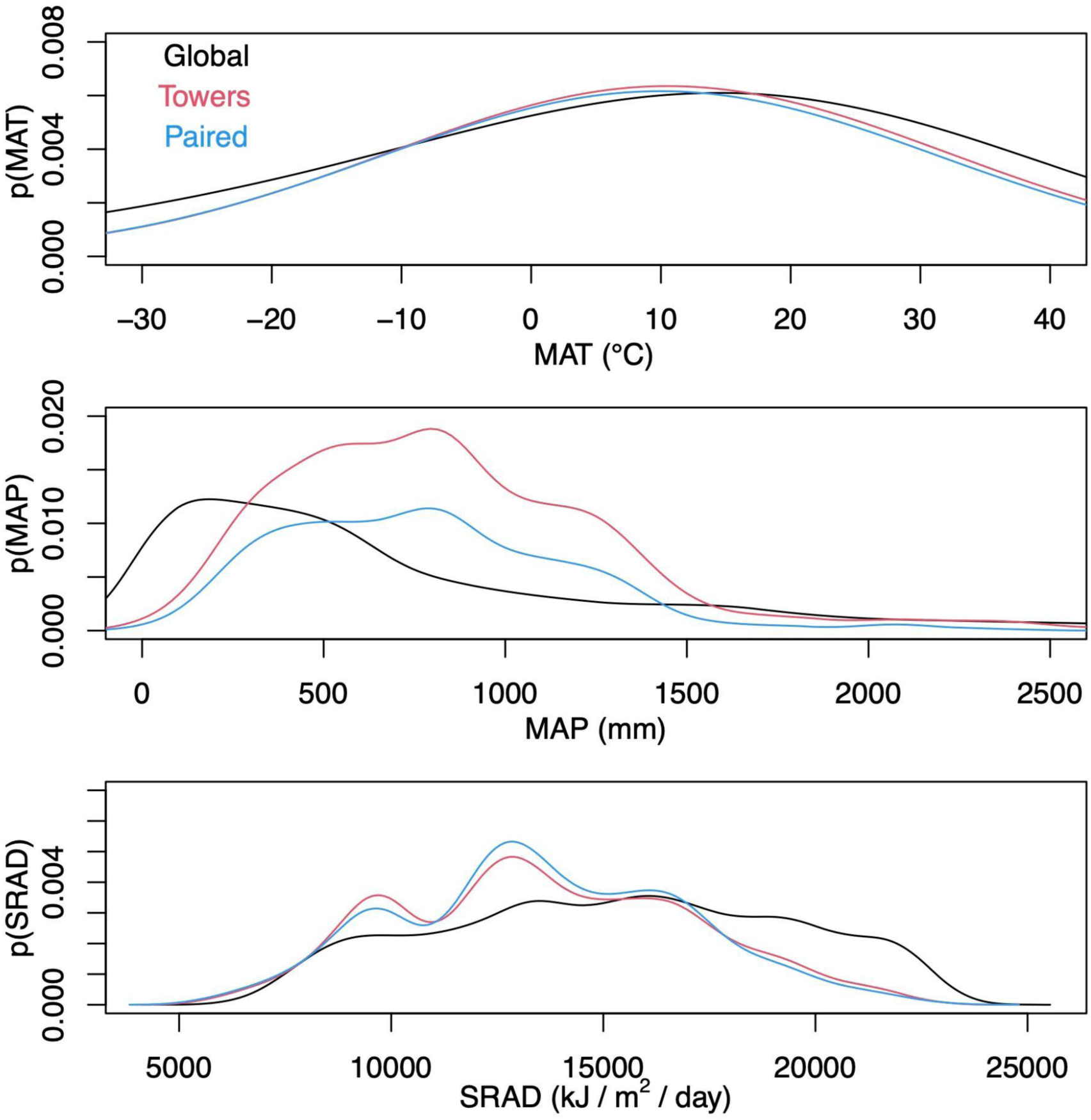
Probability density functions of mean annual temperature (MAT), mean annual precipitation (MAP) and the average daily sum of incident solar radiation (SRAD) for a random sample of 10,000 points on the terrestrial surface from WorldClim 2.1 (Fick and Hijmans, 2017) and eddy covariance tower locations for the entire database (‘Towers’, Appendix B), and those towers determined to be paired (‘Paired’, Appendix A).

**Table 1:** The percent of the terrestrial surface and eddy covariance networks with different IGBP vegetation types following Loveland et al. (2000). The total number of eddy covariance towers is listed in parentheses. Inland water bodies do not cleanly map onto IGBP vegetation types and comprise some 359 million ha globally (Bastin et al., 2019).

| IGBP<br>vegetation type | Globe | All towers | Paired towers |
| --- | --- | --- | --- |
| ENF | 4.3% | 14.4% (178) | 15.3% (106) |
| EBF | 8.3% | 4.9% (60) | 3.3 % (23) |
| DNF | 1.3% | 1.5% (19) | 1.0% (7) |
| DBF | 2.2% | 9.3% (115) | 8.1% (56) |
| MF | 4.3% | 3.1% (38) | 1.4% (10) |
| CSH | 1.8% | 1.5% (18) | 1.7% (12) |
| OSH | 12.4% | 4.5% (55) | 5.8% (40) |
| WSA | 7.0% | 1.1% (14) | 0.9% (6) |
| SAV | 6.4% | 2.6% (32) | 2.6% (18) |
| GRA | 7.6% | 16.5% (203) | 17.8% (123) |
| WET | 0.9% | 13.7% (169) | 13.9% (96) |
| CRO | 19.2% | 20.4% (251) | 22.5% (156) |
| URB | 0.2% | 2.8% (34) | 2.6% (18) |
| SNO | 11.3% | 0.2% (3) | 0.1% (1) |
| BSV | 12.7% | 1.4% (17) | 1.9% (13) |
| Water | - | 1.2% (15) | 1.0% (7) |

**Table 2:**
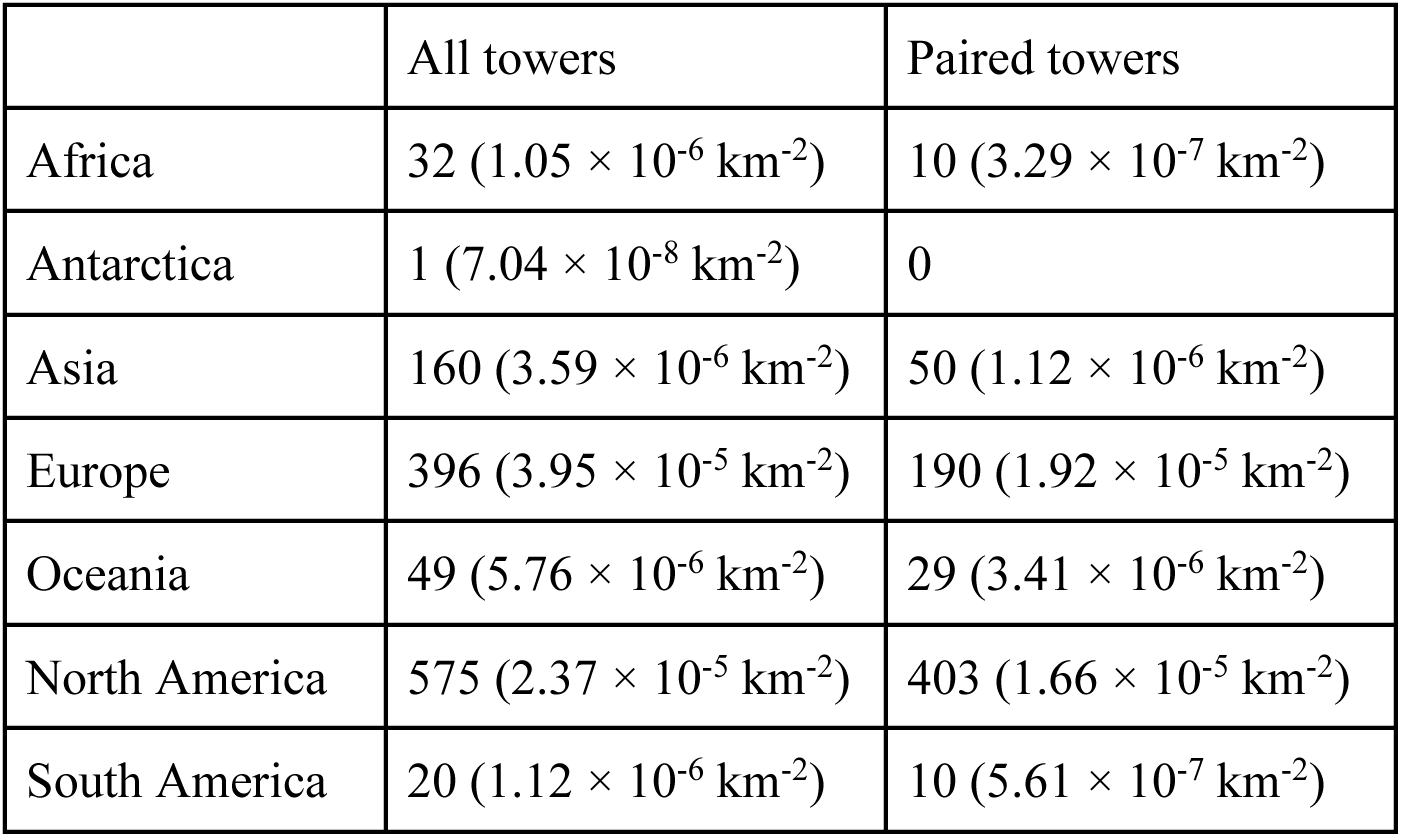
The number (density) of eddy covariance towers included in current global databases by continent.

### Climatic variables

We can address this question using the D_KL_, starting with climate. The statistical distribution of all mean annual climatic variables in the paired tower subset is more different to the global distribution of these quantities than the eddy covariance network as a whole as identified by greater D_KL_ values (Table 3). The average MAT sampled by the paired network (9.9 ℃) was not significantly different than the *unpaired* network, but warmer than the terrestrial surface (*p* = 1.7 × 10^-5^) (Tables 4 and 5). MAP in the paired network (806 mm) is closer to the global mean (739 mm) than is the full eddy covariance network (888 mm) and differences in MAP among global, paired network, and unpaired networks are all significant (Tables 4 and 5). Interestingly, the MAT and MAP for several towers exceeded the global distribution (Figure 3) which could be due to errors in reporting, or incomplete synthesis of global MAT and MAP distributions in WorldClim 2.1 as discussed in more detail below. Average SRAD across all eddy covariance towers and paired towers is less than the global mean by over 10% (Table 4, Figure 4).

**Table 3:** The Kullback-Leibler divergence (D_KL_) between global distributions of climatic and soils variables and those from the locations of all study eddy covariance towers and paired towers. The global distribution of climatic (edaphic) variables was taken from the WorldClim and the global distribution of soil texture was generated by drawing 10,000 random samples from the SoilGrids database (Poggio et al., 2021).

| Variable | All towers | Paired towers |
| --- | --- | --- |
| Elevation (m) | 0.61 | 0.61 |
| Temperature ($^{\circ}\text{C}$) | 0.60 | 0.29 |
| Precipitation (mm) | 0.46 | 0.93 |
| Solar radiation ($\text{kJ m}^{-2} \text{day}^{-1}$) | 0.20 | 0.23 |
| Sand (%) | 0.11 | 0.14 |
| Silt (%) | 0.16 | 0.18 |
| Clay (%) | 0.10 | 0.15 |
| Bulk density ($\text{g cm}^{-3}$) | 0.07 | 0.08 |
| Cation exchange capacity ($\text{cmol(c) kg}^{-1}$) | 0.13 | 0.15 |
| Soil nitrogen ($\text{g kg}^{-1}$) | 0.30 | 0.30 |
| pH | 0.27 | 0.29 |
| Soil organic carbon content ($\text{g kg}^{-1}$) | 0.26 | 0.28 |

**Table 4:** The mean (±standard deviation) value of global terrestrial climatic and edaphic variables versus those from all and paired eddy covariance towers. The global distribution of climatic variables was taken by drawing 10,000 random samples on the sphere from the WorldClim 2.1 database (Fick and Hijmans, 2017). The global distribution of soil texture was taken by drawing 10,000 random samples on the sphere from the SoilGrids 2.0 database (Poggio et al., 2021). Values for unpaired towers are presented for completeness.

| Variable | Globe | All towers | Paired towers | Unpaired towers |
| --- | --- | --- | --- | --- |
| Elevation (m) | 645 ± 839 | 433 ± 597 | 432 ± 598 | 447 ± 621 |
| Temperature ($^{\circ}$C) | 8.9 ± 18.5 | 10.5 ± 8.3 | 9.9 ± 8.1 | 11.3 ± 8.6 |
| Precipitation (mm) | 739 ± 737 | 888 ± 616 | 806 ± 479 | 987 ± 739 |
| Solar radiation ($\text{kJ m}^{-2} \text{ day}^{-1}$) | 15241 ± 4076 | 13630 ± 3430 | 13604 ± 3323 | 13623 ± 3530 |
| Sand (%) | 44.7 ± 14.0 | 41.5 ± 18.4 | 41.2 ± 19.3 | 41.6 ± 17.1 |
| Silt (%) | 30.5 ± 10.8 | 36.0 ± 12.8 | 36.4 ± 13.1 | 35.5 ± 12.4 |
| Clay (%) | 24.8 ± 7.9 | 22.5 ± 9.7 | 22.4 ± 10.4 | 22.7 ± 8.7 |
| Bulk density ($\text{g cm}^{-3}$) | 1.2 ± 0.003 | 1.1 ± 0.003 | 1.1 ± 0.003 | 1.1 ± 0.003 |
| Cation exchange capacity ($\text{cmol(c) kg}^{-1}$) | 25.2 ± 12.3 | 28.5 ± 11.8 | 28.5 ± 12.2 | 28.5 ± 11.3 |
| Soil nitrogen ($\text{g kg}^{-1}$) | 0.38 ± 0.32 | 0.58 ± 0.35 | 0.57 ± 0.35 | 0.59 ± 0.34 |
| pH | 6.4 ± 1.2 | 6.0 ± 0.9 | 6.1 ± 0.9 | 5.9 ± 0.96 |
| Soil organic carbon content ($\text{g kg}^{-1}$) | 5.4 ± 5.9 | 8.3 ± 7.3 | 8.1 ± 7.6 | 8.6 ± 7.0 |

**Table 5:**
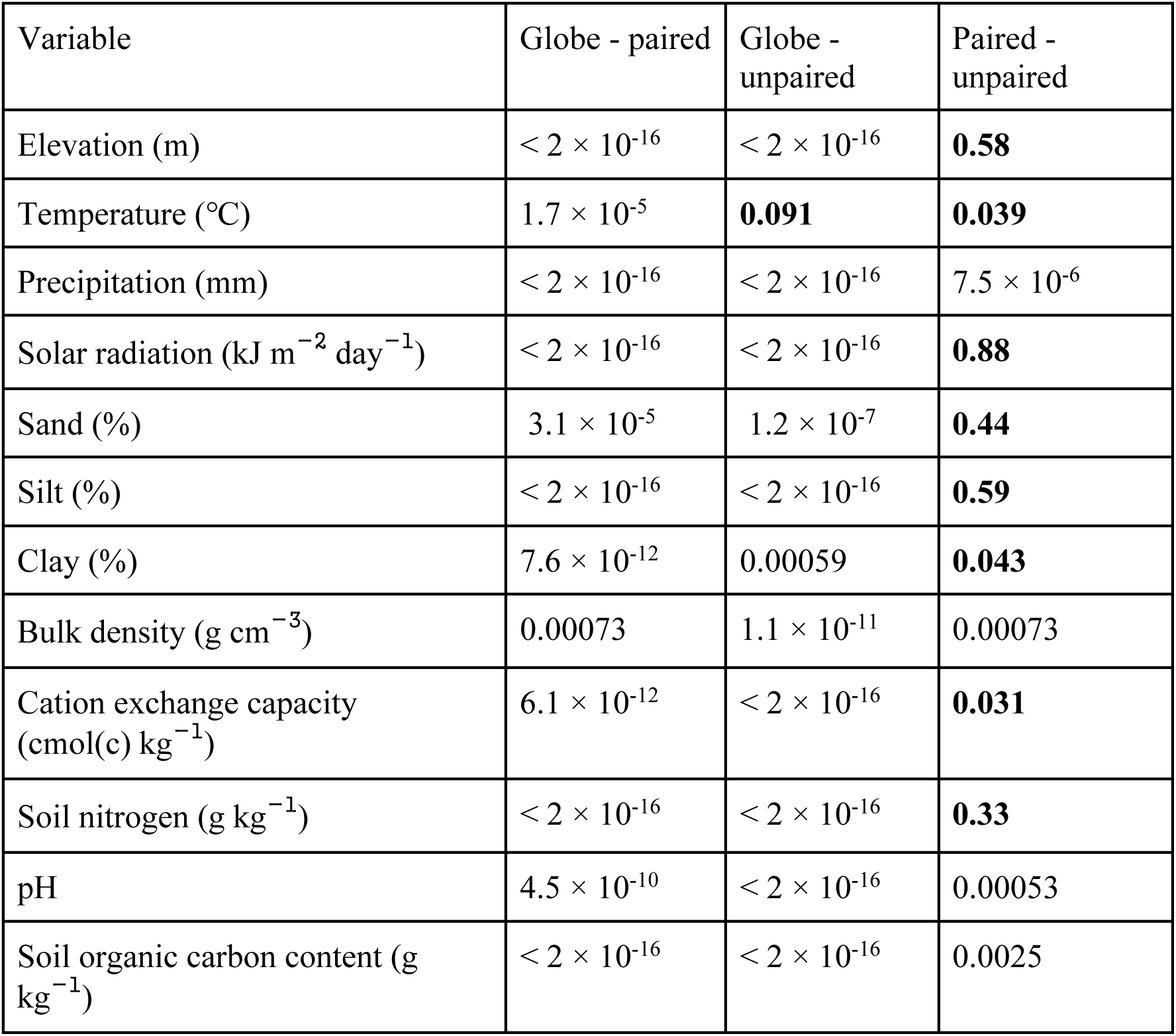
The likelihood of significant differences of the mean (*p*-values) between random draws of climatic and edaphic variables (‘Globe’), the paired tower network, and the unpaired tower network (i.e. all towers excluding paired towers) from pairwise comparisons using the Wilcoxon rank sum test with continuity and Holm-Bonferroni corrections. Values that are not significantly different at the α < 0.01 level are indicated in bold.

### Edaphic variables

The paired tower network captures a similar distribution of soil textures to the *unpaired* eddy covariance network, but this differs from the rest of the terrestrial surface (Tables 4&5, Figures 5&6). Mean clay content in paired towers (22.4%) is about 10% less than the global mean (24.8%), but global terrestrial soils have some 8% more sand and nearly 20% less silt, on average, compared to paired towers and the entire tower database, and these differences are statistically significant (Tables 4&5). Differences in soil texture are reflected in the soil texture triangles (Figure 5), and the distributions of each soil texture component (Figure 6) further reveal that the tower networks have a higher mode in average silt content, which points to the suggestion that global towers may sample more fertile soils than the global terrestrial soils on average.

**Figure 5:**
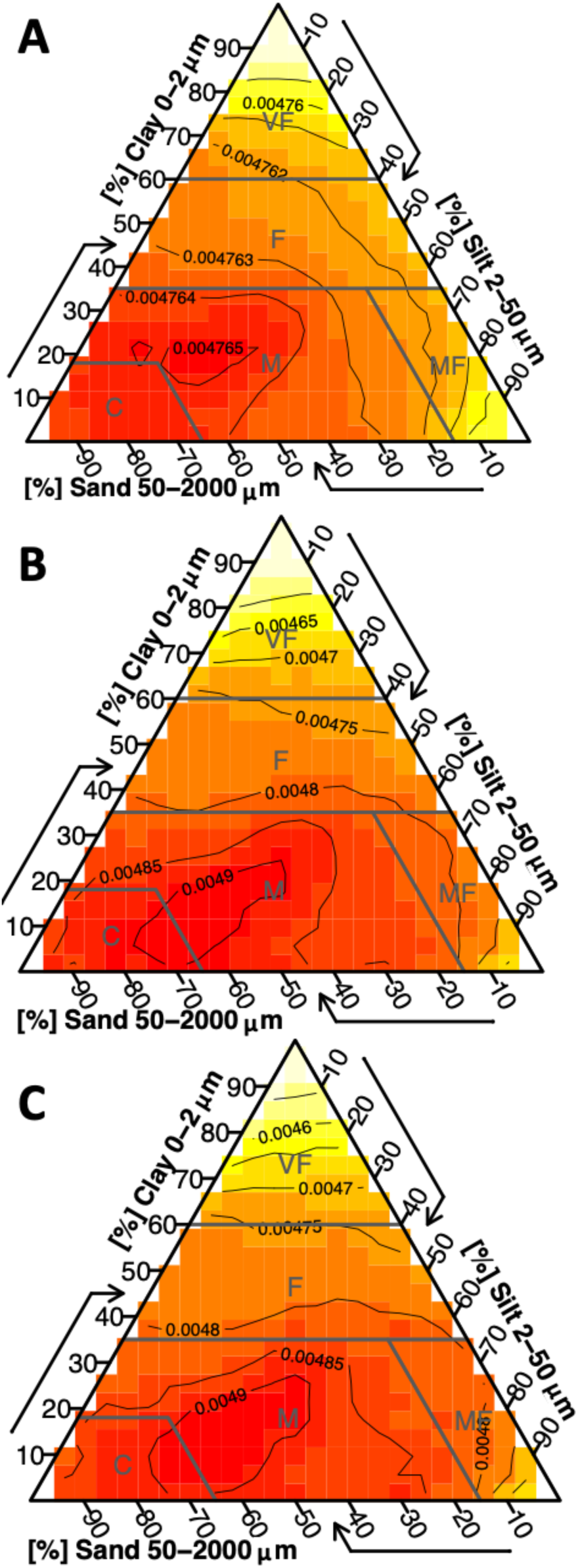
A heat map of the global distribution of soil texture generated by drawing (A) 10,000 random samples, (B) data from all eddy covariance towers, and (C) data from paired eddy covariance towers from the SoilGrids database (Poggio et al., 2021). All figures were generated using the *soiltexture* package (Moeys, 2018) in R.

**Figure 6:**
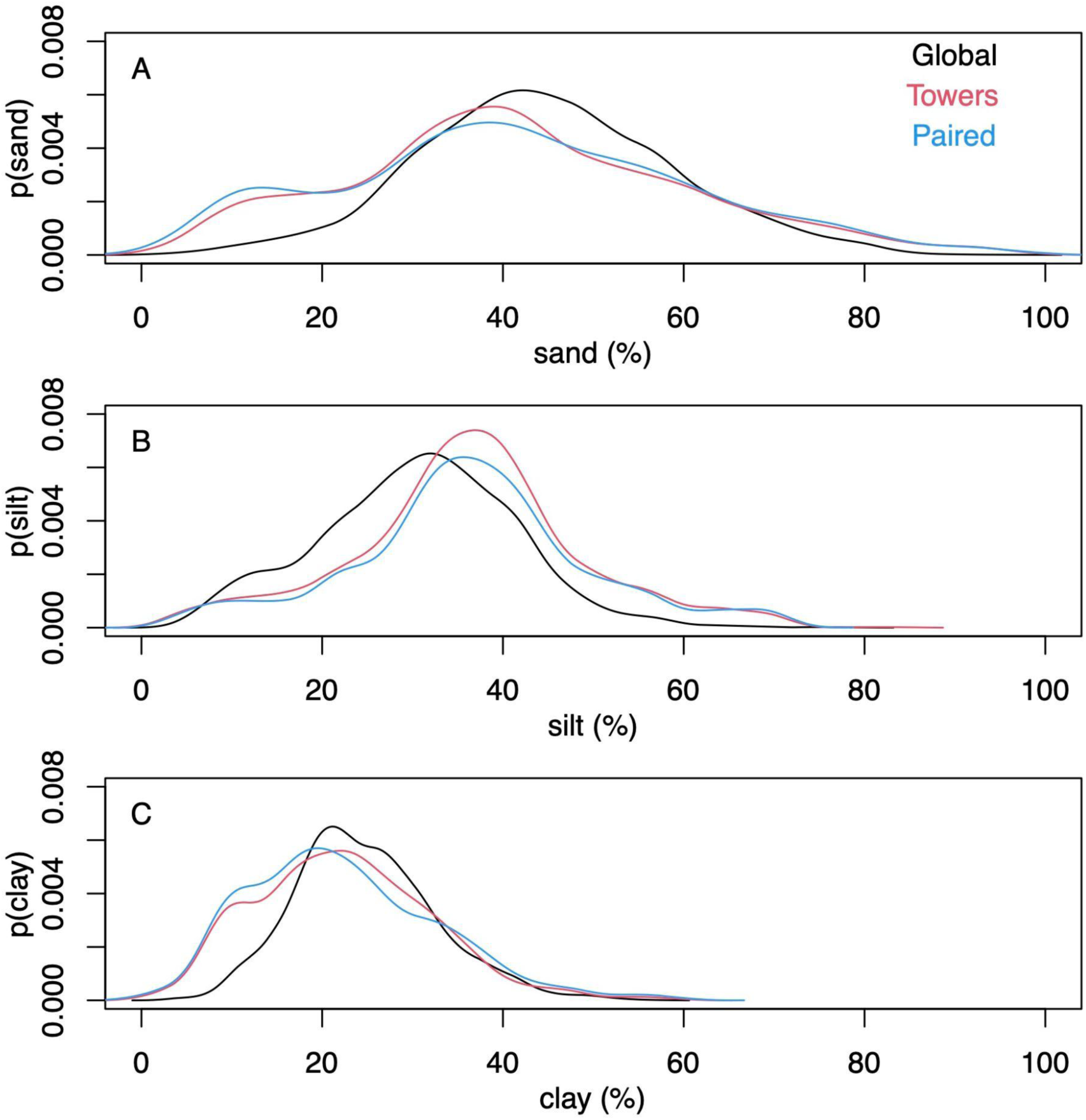
Probability density functions of sand (A), silt (B), and clay (C) fractions for a random sample of 10,000 points on the terrestrial surface from SoilGrids 2.0 (Poggio et al., 2021) and eddy covariance tower locations for the entire database (‘Towers’, Appendix B), and those towers determined to be paired (‘Paired’, Appendix A).

Soil N in the upper 5 cm is ∼50% greater on average across the tower network (0.58 g/kg) and paired towers (0.57 g/kg) than the terrestrial land surface (0.38 g/kg) (Table 3, Figure 7). SOC is also greater in the eddy covariance network (8.3 g/kg) and paired towers (8.1 g/kg) than terrestrial soils on average (5.4 g/kg), and average cation exchange capacity is over 10% greater in tower locations than terrestrial soils on average (Table 3, Figure 7). Average soil pH is slightly more acidic across eddy covariance sites (mean pH ∼ 6) than the average value for the terrestrial surface (pH = 6.4) due in part to an undersampling of basic soils with a secondary peak of pH around 8 (Figure 7). Combined with soil textural observations, soil chemistry and biogeochemistry measurements suggest that the eddy covariance network samples soils that are more fertile than the entire terrestrial surface, on average, although the paired tower and unpaired flux databases sample a similar distribution of soil characteristics (Tables 4 and 5).

**Figure 7:**
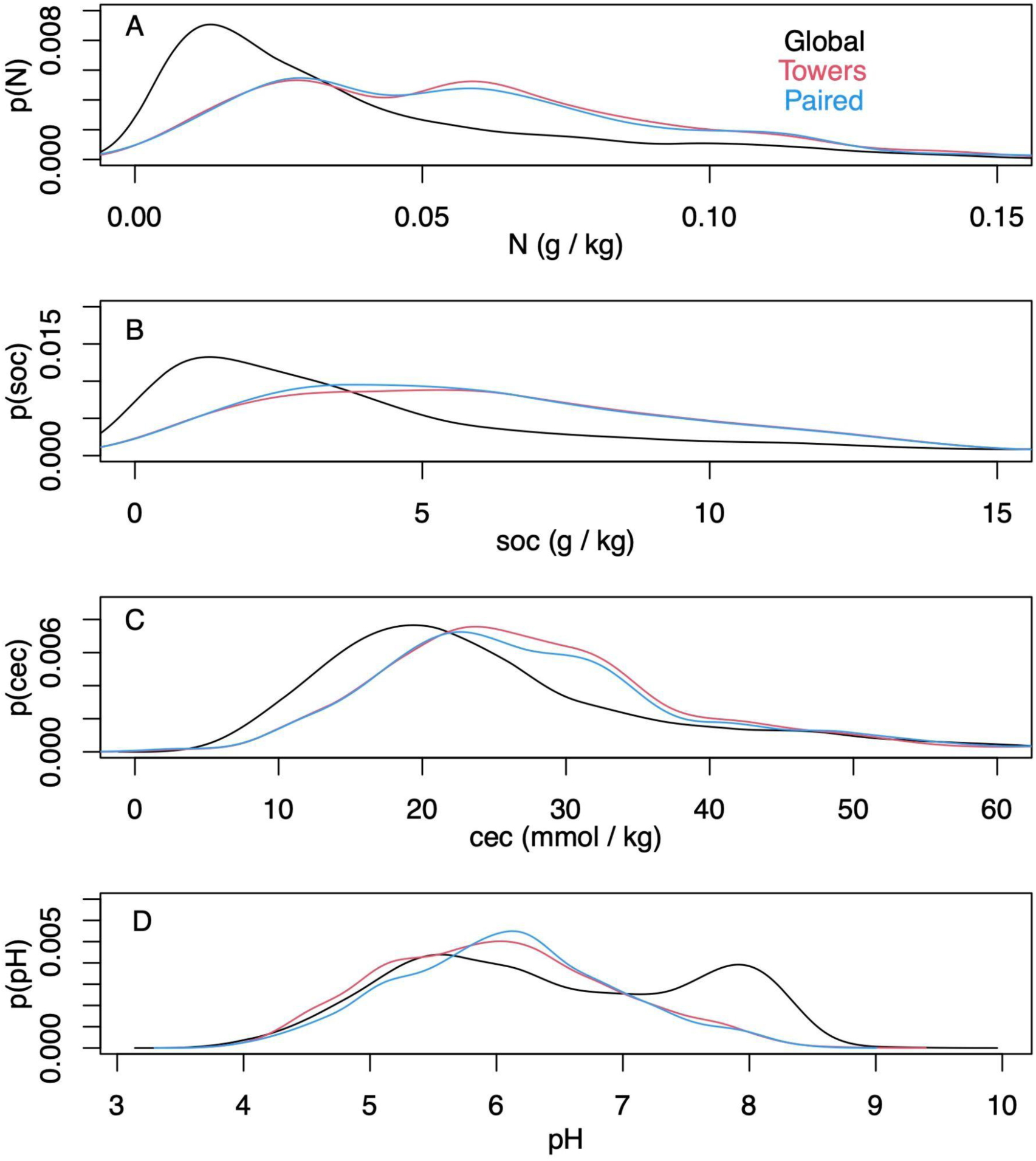
Probability density functions of soil nitrogen (A), soil organic carbon (soc, B), cation exchange capacity (cec, C), and pH (D) for a random sample of 10,000 points on the terrestrial surface from SoilGrids2.0 (Poggio et al., 2021) and eddy covariance tower locations for the entire database (‘Towers’, Appendix B), and those towers determined to be paired (‘Paired’, Appendix A).

These soil characteristics can vary considerably across space and across the different datasets used to infer them. For example, across the 14 eddy covariance towers in the CHEESEHEAD19 domain from which soil texture data from cores, the USGS/NRCS database, and the SoilGrids2.0 database were all available (Table 6), SoilGrids2.0 had lower sand content (60.7% versus ∼75% for the other datasets), higher clay content (9.6% versus ∼ 6%), and higher silt content (29.8% versus ∼19-20%). This is not to say that the soil core and USGS data matched across each individual site: these differed by over 25% (for the case of sand at the SE2 site). In brief, one can expect uncertainties when estimating difficult-to-measure edaphic variables that often exhibit considerable heterogeneity (Mulla and McBratney, 2002); note for example that the USGS/NRCS database estimates sand content that ranges from 92.7% at NE2 to 49.5% at SE2, a difference of over 40% within the 10 × 10 km CHEESEHEAD19 domain.

**Table 6:** Soil texture characteristics from the CHEESEHEAD19 experiment (Butterworth et al., 2021) inferred from soil cores within the eddy covariance flux footprint, the USGS/NRCS database at the flux tower locations (Shveytser et al., 2022), and the SoilGrids 2.0 database (Poggio et al., 2021) at the flux tower locations.

|  | Cores |  |  | USGS |  |  | SoilGrids |  |  |
| --- | --- | --- | --- | --- | --- | --- | --- | --- | --- |
| Site Name | sand | silt | clay | sand | silt | clay | sand | silt | clay |
| NW1 | 83.0 | 11.8 | 5.2 | 68.4 | 21.1 | 10.5 | 57 | 32.8 | 10.2 |
| NW2 | 77.3 | 14.3 | 8.4 | 65.1 | 29.5 | 5.4 | 57.7 | 31.7 | 10.6 |
| NW3 | NA | NA | NA | 56.4 | 32.7 | 10.9 | 56.7 | 32.2 | 11.1 |
| NW4 | NA | NA | NA | 56.4 | 32.7 | 10.9 | 56.1 | 33.2 | 10.7 |
| NE1 | 77.2 | 16.5 | 6.3 | 90.2 | 6.6 | 3.2 | 60 | 29.2 | 10.8 |
| NE2 | 79.9 | 15.1 | 5.0 | 92.7 | 3.6 | 3.7 | 62.1 | 29.3 | 8.6 |
| NE3 | 77.3 | 15.8 | 6.9 | 65.1 | 29.5 | 5.4 | 59.1 | 32.4 | 8.5 |
| NE4 | 72.3 | 20.1 | 7.6 | 65.1 | 29.5 | 5.4 | 61.9 | 29.8 | 8.2 |
| SW1 | 71.6 | 21.2 | 7.2 | 65.1 | 29.5 | 5.4 | 59.7 | 28.5 | 11.9 |
| SW2 | 77.1 | 19.2 | 3.6 | 90.2 | 6.6 | 3.2 | 56.9 | 31.1 | 12 |
| SW3 | 84.2 | 13.6 | 2.2 | 65.1 | 29.5 | 5.4 | 57.9 | 32 | 10.1 |
| SW4 | 66.7 | 28.2 | 5.1 | 65.1 | 29.5 | 5.4 | 59.3 | 31.9 | 8.8 |
| SE2 | 77.0 | 19.8 | 3.1 | 49.5 | 42.2 | 8.3 | 63.9 | 27.3 | 8.8 |
| SE3 | 67.1 | 25.5 | 7.4 | 79.1 | 12.4 | 8.5 | 63.4 | 28.2 | 8.4 |
| SE4 | NA | NA | NA | 56.4 | 32.7 | 10.9 | 62.9 | 28.6 | 8.6 |
| SE5 | 71.1 | 21.1 | 7.8 | 90.2 | 6.6 | 3.2 | 66.6 | 25.4 | 7.9 |
| SE6 | 71.3 | 22.3 | 6.4 | 90.2 | 6.6 | 3.2 | 63.8 | 27.2 | 9 |

Further to the point of data uncertainty, the mean (± standard deviation) difference between tower reported and WorldClim 2.1 MAT values is 0.27 ± 2.29 ℃ with a maximum difference, somehow, of 32 ℃. The mean difference in MAP between tower reported and WorldClim 2.1 information is 28.3 ± 361 mm with a maximum difference of, remarkably, 8678 mm. These differences arise for multiple reasons including clerical errors when creating eddy covariance metadata, uncertainties in reporting, uncertainties in tower measurements, and uncertainties in climate datasets.

## Discussion

Paired eddy covariance towers sample more biased distributions of climate space than the eddy covariance network alone, as we anticipated (Table 3). We did not expect the paired towers to sample a similar edaphic space to the entire eddy covariance network as we observed, but both networks tended to sample global richer soils (e.g., with more N, SOC, and silt) than the global average (Tables 5&6). There are key geographic discrepancies in the eddy covariance network and paired tower network (Chu et al., 2017; Pastorello et al., 2020; Xiao et al., 2012), which tend to sample ecosystems in Europe, Australia, and North America (Table 2) with fewer opportunities to compare ecosystems in tropical, dry, and cold climates (Table 1; Figures 2-4).

More opportunities to use the paired tower approach exist now that the NEON, AmeriFlux, ICOS, TERN and other measurement networks provide baseline eddy covariance data from well-studied ecosystems to which additional towers can be paired. As currently operating, these networks will not resolve global discrepancies in the geographic distribution of flux towers or paired flux towers (e.g., Figure 2) as they mostly add observational capacity to the United States, Europe, and Australia.

These discrepancies should be addressed, especially in light of the growing interest in nature-based solutions to mitigate the effects of and to adapt to climate change (Hemes et al., 2021; Seddon, 2022), which emphasizes the need to understand trace gas, water, and energy fluxes across global ecosystems that will be essential for quantifying baseline, additionality, and permanence metrics required of carbon credit markets. Paired sites can help design effective, defensible climate solutions and policies (Novick, 2022), but at the present their scope is limited to certain regions, whereas global change is of global concern.

### Replicability

A major criticism of eddy covariance studies is that they are rarely ‘replicated’ despite the major expense that this would entail or the likelihood that the measurements would likely be pseudoreplicated if the measurements are not in ecosystems that are ‘independent’ regardless of the independence of flux footprints themselves. It has been argued that multiple flux towers are necessary to fully sample a given ecosystem (Hill et al., 2016), because fluxes from different parts of the same ecosystem may differ due to vegetative or edaphic characteristics (Oren et al., 2006). This raises the question of whether multiple towers in a single ecosystem are ‘paired’. By our definition we argue that they would be, as they are inevitably capturing a different distribution of vegetative or edaphic characteristics within their flux footprint. A more relevant question may be if models or remote sensing algorithms can adequately capture the spatial and temporal variability in fluxes that may exist within what we may consider a single ‘ecosystem’, and if the representativeness of tower observations across space (Chu et al., 2021) and time (Chu et al., 2017) make it possible to link them to larger scales. In other words, lessons can be learned from multiple towers located in a single ecosystem that can help us understand the rich diversity of fluxes that can arise from spatially and temporally heterogeneous ecosystem function.

### Uncertainties

Climate changes dynamically across time and space. The use of MAT and MAP in our analysis therefore introduces a shifting baseline as climate continues to change, but we know of no way to encapsulate the distribution of climatic variables that is more commonly used or straightforward. These variables are prone to reporting uncertainty as suggested by Figure 3 and the comparison of MAT and MAP values in the *Results* where it was noted that MAT (MAP) differences between tower reporting and global databases differ by up to 32 ℃ (8678 mm). From this analysis it should be clear that global climate datasets reasonably simulate tower climatic conditions in many instances, but both tower and climate dataset information should be double-checked for logic when reporting. We use reported values here to help quantify the degree of uncertainty that climate reporting and extrapolation can introduce. Uncertainties in climate data reporting are unlikely to impact our overarching results as it is clear from the global distribution that fewer towers exist in regions both hot and cold, and wet and dry (Figures 2-3).

The challenge of simplifying edaphic characteristics is less straightforward as these vary – often dramatically – across space and soil depth, and sometimes across time in the case of extreme events or management, all of which impact how we interpret eddy covariance observations (Herbst et al., 2021; Levy et al., 2020; Oren et al., 2006; Rey-Sanchez et al., 2022; Tuovinen et al., 2019; Vanderborght et al., 2010). Data products like SoilGrids 2.1 represent the state-of-the-art of our understanding of the global distribution of soil characteristics but from our analysis substantial uncertainties may result when applying such data to individual tower locations (Table 6). Whereas such uncertainties inevitably impact our complication of edaphic characteristics at each tower (Appendices A&B) and therefore need to be considered when interpreting results, we feel that the overarching result that eddy covariance towers sample regions of the globe with richer soils is largely robust. The lack of measurement towers across deserts and other dryland regions and a disproportionate number of towers in croplands (which, admittedly, are disproportionately important for human health and wellbeing) suggest that the full distribution of soil fertility is poorly captured. Global analyses should be cognizant of this edaphic bias. We note that there is a large undersampling of ecosystems with acidic soils, especially those with pH ∼ 8 that are buffered by calcium carbonate, leaving opportunities to study abrupt spatial heterogeneity in soil characteristics including pH that are largely controlled by aridity (Slessarev et al., 2016).

### Metadata and information availability

It is apparent from Appendices A and B that key information is unavailable for some sites, which stands to reason if tower data and metadata are often shared on a volunteer basis (Novick et al., 2018) that may have no mandate or additional funding to support data sharing. From this perspective the FLUXNET network and regional networks are exemplary in data dissemination, but key pieces of information for a paired tower synthesis were difficult to glean. These include the intent of the towers (e.g., whether they were intentionally paired or not) and the specific objectives that studies were designed to address or ecosystem transitions they were designed to measure. A formal meta-analysis of paired tower data would require such information, which would be an undertaking but would lead to remarkable insights given the key findings of paired tower syntheses to date (e.g. Luyssaert et al., 2014; Zhang et al., 2020). We advocate for additional information sharing for datasets for which sharing is allowed and look forward to new FLUXNET synthetic activities that build off of the legacy of the Marconi (Falge et al., 2006), La Thuile (Verma et al., 2014), and FLUXNET2015 (Pastorello et al., 2020) datasets.

## Conclusions

Individual and paired eddy covariance studies arise to address pressing questions in surface-atmosphere exchange. Only recently have towers been sited at locations with large-scale representativeness in mind (Schimel et al., 2007), so one should not expect that towers across the globe have representative distributions across climatic and edaphic space. Instead, this is a goal to which we should aspire for equity in ecosystem-level measurements that are important in the global climate system. The current tower bias toward temperate regions – especially for paired towers – leaves opportunities to expand the flux network globally and further build international partnerships to help do so. Our analysis also demonstrates that current flux tower placement samples soils with richer characteristics than the global average but the entire tower network and paired towers sample similar edaphic variability, on average. Additional information about tower measurements and their intent will be useful for future studies of paired tower findings, all of which will continue to benefit from the remarkable energy and enthusiasm that the eddy covariance community has used to create our state-of-the-art observatory of surface-atmosphere exchange.

## Supporting information

Appendix B, See also https://figshare.com/articles/dataset/Location_vegetation_climate_and_soils_information_from_global_paired_eddy_covariance_towers

Appendix A, See also https://figshare.com/articles/dataset/Location_vegetation_climate_and_soils_information_from_global_paired_eddy_covariance_towers

## Acknowledgements

First and foremost, we are grateful to the global eddy covariance community for measuring fluxes and sharing data, and regional networks like FLUXNET, AmeriFlux, ICOS, European Eddy Fluxes Database, OzFlux, AsiaFlux for hosting and maintaining the databases. We acknowledge support from the U.S. National Science Foundation Macrosystems Biology award 2106012 and the University of Wisconsin Research Forward program. Hannah Koch, Sophie Hoffman, and others provided insightful comments for which we are grateful. Jamie Cleverly and Peter Issac provided key insights into eddy covariance sites in Australia and New Zealand.

## Appendices

Appendices A and B are publicly available at https://doi.org/10.6084/m9.figshare.22189693.v3

## References

1. Afgani, M., Sinanovic, S., Haas, H., 2008. Anomaly detection using the Kullback-Leibler divergence metric. Presented at the 2008 First International Symposium on Applied Sciences on Biomedical and Communication Technologies, IEEE, pp. 1–5.

2. Amiro, B.D., 2001. Paired-tower measurements of carbon and energy fluxes following disturbance in the boreal forest. Glob. Change Biol. 7, 253–268.

3. Anapalli, S.S., Fisher, D.K., Reddy, K.N., Krutz, J.L., Pinnamaneni, S.R., Sui, R., 2019. Quantifying water and CO2 fluxes and water use efficiencies across irrigated C3 and C4 crops in a humid climate. Sci. Total Environ. 663, 338–350.

4. Anderson, R.G., Wang, D., 2014. Energy budget closure observed in paired Eddy Covariance towers with increased and continuous daily turbulence. Agric. For. Meteorol. 184, 204–209.

5. Baddeley, A., Lawrence, T., Rubak, E., 2017. globe: Plot 2D and 3D Views of the Earth, Including Major Coastline.

6. Baker, J., Griffis, T., 2005. Examining strategies to improve the carbon balance of corn/soybean agriculture using eddy covariance and mass balance techniques. Agric. For. Meteorol. 128, 163–177.

7. Baker, T.P., Jordan, G.J., Steel, E.A., Fountain-Jones, N.M., Wardlaw, T.J., Baker, S.C., 2014. Microclimate through space and time: microclimatic variation at the edge of regeneration forests over daily, yearly and decadal time scales. For. Ecol. Manag. 334, 174–184.

8. Baldocchi, D., 2014. Measuring fluxes of trace gases and energy between ecosystems and the atmosphere–the state and future of the eddy covariance method. Glob. Change Biol. 20, 3600–3609.

9. Baldocchi, D., 2008. ‘Breathing’ of the terrestrial biosphere: lessons learned from a global network of carbon dioxide flux measurement systems. Aust. J. Bot. 56, 1–26.

10. Baldocchi, D., Ma, S., 2013. How will land use affect air temperature in the surface boundary layer? Lessons learned from a comparative study on the energy balance of an oak savanna and annual grassland in California, USA. Tellus B Chem. Phys. Meteorol. 65, 19994.

11. Baldocchi, D.D., 2020. How eddy covariance flux measurements have contributed to our understanding of Global Change Biology. Glob. Change Biol. 26, 242–260.

12. Barr, A., Morgenstern, K., Black, T., McCaughey, J., Nesic, Z., 2006. Surface energy balance closure by the eddy-covariance method above three boreal forest stands and implications for the measurement of the CO_2_ flux. Agric. For. Meteorol. 140, 322–337.

13. Bastin, L., Gorelick, N., Saura, S., Bertzky, B., Dubois, G., Fortin, M.-J., Pekel, J.-F., 2019. Inland surface waters in protected areas globally: Current coverage and 30-year trends. PloS One 14, e0210496.

14. Bavin, T., Griffis, T., Baker, J., Venterea, R., 2009. Impact of reduced tillage and cover cropping on the greenhouse gas budget of a maize/soybean rotation ecosystem. Agric. Ecosyst. Environ. 134, 234–242.

15. Beaudette, D., Skovlin, J., Roecker, S., Beaudette, M.D., 2022. Package ‘soilDB.’

16. Beringer, J., Hutley, L.B., McHugh, I., Arndt, S.K., Campbell, D., Cleugh, H.A., Cleverly, J., Resco de Dios, V., Eamus, D., Evans, B., 2016. An introduction to the Australian and New Zealand flux tower network–OzFlux. Biogeosciences 13, 5895–5916.

17. Burakowski, E., Tawfik, A., Ouimette, A., Lepine, L., Novick, K., Ollinger, S., Zarzycki, C., Bonan, G., 2018. The role of surface roughness, albedo, and Bowen ratio on ecosystem energy balance in the Eastern United States. Agric. For. Meteorol. 249, 367–376.

18. Butterworth, B.J., Desai, A.R., Townsend, P.A., Petty, G.W., Andresen, C.G., Bertram, T.H., Kruger, E.L., Mineau, J.K., Olson, E.R., Paleri, S., 2021. Connecting land–atmosphere interactions to surface heterogeneity in CHEESEHEAD19. Bull. Am. Meteorol. Soc. 102, E421–E445.

19. Chi, J., Waldo, S., Pressley, S., O’Keeffe, P., Huggins, D., Stöckle, C., Pan, W.L., Brooks, E., Lamb, B., 2016. Assessing carbon and water dynamics of no-till and conventional tillage cropping systems in the inland Pacific Northwest US using the eddy covariance method. Agric. For. Meteorol. 218, 37–49.

20. Chu, H., Baldocchi, D.D., John, R., Wolf, S., Reichstein, M., 2017. Fluxes all of the time? A primer on the temporal representativeness of FLUXNET. J. Geophys. Res. Biogeosciences 122, 289–307.

21. Chu, H., Luo, X., Ouyang, Z., Chan, W.S., Dengel, S., Biraud, S.C., Torn, M.S., Metzger, S., Kumar, J., Arain, M.A., 2021. Representativeness of Eddy-Covariance flux footprints for areas surrounding AmeriFlux sites. Agric. For. Meteorol. 301, 108350.

22. Cleverly, J., Vote, C., Isaac, P., Ewenz, C., Harahap, M., Beringer, J., Campbell, D.I., Daly, E., Eamus, D., He, L., 2020. Carbon, water and energy fluxes in agricultural systems of Australia and New Zealand. Agric. For. Meteorol. 287, 107934.

23. Clifton, O.E., Fiore, A.M., Massman, W.J., Baublitz, C.B., Coyle, M., Emberson, L., Fares, S., Farmer, D.K., Gentine, P., Gerosa, G., 2020. Dry deposition of ozone over land: processes, measurement, and modeling. Rev. Geophys. 58, e2019RG000670.

24. Delwiche, K.B., Knox, S.H., Malhotra, A., Fluet-Chouinard, E., McNicol, G., Feron, S., Ouyang, Z., Papale, D., Trotta, C., Canfora, E., 2021. FLUXNET-CH_4_: a global, multi-ecosystem dataset and analysis of methane seasonality from freshwater wetlands. Earth Syst. Sci. Data 13, 3607–3689.

25. Desai, A.R., Bolstad, P.V., Cook, B.D., Davis, K.J., Carey, E.V., 2005. Comparing net ecosystem exchange of carbon dioxide between an old-growth and mature forest in the upper Midwest, USA. Agric. For. Meteorol. 128, 33–55.

26. Deshmukh, C.S., Julius, D., Evans, C.D., Susanto, A.P., Page, S.E., Gauci, V., Laurén, A., Sabiham, S., Agus, F., Asyhari, A., 2020. Impact of forest plantation on methane emissions from tropical peatland. Glob. Change Biol. 26, 2477–2495.

27. Detto, M., Katul, G.G., Siqueira, M., Juang, J.-Y., Stoy, P., 2008. The structure of turbulence near a tall forest edge: The backward-facing step flow analogy revisited. Ecol. Appl. 18, 1420–1435.

28. Duman, T., Huang, C.-W., Litvak, M.E., 2021. Recent land cover changes in the Southwestern US lead to an increase in surface temperature. Agric. For. Meteorol. 297, 108246.

29. Eder, F., Schmidt, M., Damian, T., Träumner, K., Mauder, M., 2015. Mesoscale eddies affect near-surface turbulent exchange: Evidence from lidar and tower measurements. J. Appl. Meteorol. Climatol. 54, 189–206.

30. Eder, F., Serafimovich, A., Foken, T., 2013. Coherent structures at a forest edge: properties, coupling and impact of secondary circulations. Bound.-Layer Meteorol. 148, 285–308.

31. Falge, E., Aubinet, M., Bakwin, P., Baldocchi, D., Berbigier, P., Bernhofer, C., Black, T., Ceulemans, R., Davis, K., Dolman, A., 2006. FLUXNET Marconi conference gap-filled flux and meteorology data, 1992-2000. ORNL DAAC.

32. Fick, S.E., Hijmans, R.J., 2017. WorldClim 2: new 1-km spatial resolution climate surfaces for global land areas. Int. J. Climatol. 37, 4302–4315.

33. French, A.N., Hunsaker, D.J., Sanchez, C.A., Saber, M., Gonzalez, J.R., Anderson, R., 2020. Satellite-based NDVI crop coefficients and evapotranspiration with eddy covariance validation for multiple durum wheat fields in the US Southwest. Agric. Water Manag. 239, 106266.

34. Goulden, M.L., Winston, G.C., McMillan, A., Litvak, M.E., Read, E.L., Rocha, A.V., Rob Elliot, J., 2006. An eddy covariance mesonet to measure the effect of forest age on land– atmosphere exchange. Glob. Change Biol. 12, 2146–2162.

35. Hargrove, W., Hoffman, F., Law, B., 2003. New analysis reveals representativeness of the AmeriFlux network. Eos Trans. Am. Geophys. Union 84, 529–535.

36. Haskett, J.D., Pachepsky, Y.A., Acock, B., 1995. Use of the beta distribution for parameterizing variability of soil properties at the regional level for crop yield estimation. Agric. Syst. 48, 73–86.

37. Heinsch, F.A., Zhao, M., Running, S.W., Kimball, J.S., Nemani, R.R., Davis, K.J., Bolstad, P.V., Cook, B.D., Desai, A.R., Ricciuto, D.M., 2006. Evaluation of remote sensing based terrestrial productivity from MODIS using regional tower eddy flux network observations. IEEE Trans. Geosci. Remote Sens. 44, 1908–1925.

38. Hemes, K.S., Chamberlain, S.D., Eichelmann, E., Anthony, T., Valach, A., Kasak, K., Szutu, D., Verfaillie, J., Silver, W.L., Baldocchi, D.D., 2019. Assessing the carbon and climate benefit of restoring degraded agricultural peat soils to managed wetlands. Agric. For. Meteorol. 268, 202–214.

39. Hemes, K.S., Runkle, B.R., Novick, K.A., Baldocchi, D.D., Field, C.B., 2021. An ecosystem-scale flux measurement strategy to assess natural climate solutions. Environ. Sci. Technol. 55, 3494–3504.

40. Herbst, M., Pohlig, P., Graf, A., Weihermüller, L., Schmidt, M., Vanderborght, J., Vereecken, H., 2021. Quantification of water stress induced within-field variability of carbon dioxide fluxes in a sugar beet stand. Agric. For. Meteorol. 297, 108242.

41. Hill, T., Chocholek, M., Clement, R., 2016. The case for increasing the statistical power of eddy covariance ecosystem studies: why, where and how? Glob. Change Biol. 23, 2154–2165.

42. Hollinger, D., Richardson, A., 2005. Uncertainty in eddy covariance measurements and its application to physiological models. Tree Physiol. 25, 873–885.

43. Hollister, J., Shah, T., Robitaille, A., Beck, M., Johnson, M., 2022. elevatr: access elevation data from various APIs. R Package Version 042 0.4.2. https://doi.org/10.5281/zenodo.5809645

44. Hu, J., Murphy, B., Huang, J., 2020. Soil Texture, vis-NIR Spectra, and Derived Soil Chemistry. Version 1.0. UCAR/NCAR - Earth Observing Laboratory.

45. Juang, J., Katul, G., Siqueira, M., Stoy, P., Novick, K., 2007a. Separating the effects of albedo from eco-physiological changes on surface temperature along a successional chronosequence in the southeastern United States. Geophys. Res. Lett. 34.

46. Juang, J., Porporato, A., Stoy, P.C., Siqueira, M.S., Oishi, A.C., Detto, M., Kim, H., Katul, G.G., 2007b. Hydrologic and atmospheric controls on initiation of convective precipitation events. Water Resour. Res. 43.

47. Juang, J.-Y., Katul, G.G., Porporato, A., Stoy, P.C., Siqueira, M.S., Detto, M., Kim, H., Oren, R., 2007. Eco-hydrological controls on summertime convective rainfall triggers. Glob. Change Biol. 13, 887–896.

48. Jung, M., Reichstein, M., Bondeau, A., 2009. Towards global empirical upscaling of FLUXNET eddy covariance observations: validation of a model tree ensemble approach using a biosphere model. Biogeosciences 6, 2001–2013.

49. Kessomkiat, W., Franssen, H.-J.H., Graf, A., Vereecken, H., 2013. Estimating random errors of eddy covariance data: An extended two-tower approach. Agric. For. Meteorol. 171, 203– 219.

50. Kowalski, A.S., Loustau, D., Berbigier, P., Manca, G., Tedeschi, V., Borghetti, M., Valentini, R., Kolari, P., Berninger, F., Rannik, Ü., 2004. Paired comparisons of carbon exchange between undisturbed and regenerating stands in four managed forests in Europe. Glob. Change Biol. 10, 1707–1723.

51. Krauss, K.W., Holm Jr, G.O., Perez, B.C., McWhorter, D.E., Cormier, N., Moss, R.F., Johnson, D.J., Neubauer, S.C., Raynie, R.C., 2016. Component greenhouse gas fluxes and radiative balance from two deltaic marshes in Louisiana: Pairing chamber techniques and eddy covariance. J. Geophys. Res. Biogeosciences 121, 1503–1521.

52. Kullback, S., Leibler, R.A., 1951. On information and sufficiency. Ann. Math. Stat. 22, 79–86.

53. Lasslop, G., Reichstein, M., Kattge, J., Papale, D., 2008. Influences of observation errors in eddy flux data on inverse model parameter estimation. Biogeosciences 5, 1311–1324.

54. Levy, P., Drewer, J., Jammet, M., Leeson, S., Friborg, T., Skiba, U., Van Oijen, M., 2020. Inference of spatial heterogeneity in surface fluxes from eddy covariance data: A case study from a subarctic mire ecosystem. Agric. For. Meteorol. 280, 107783.

55. Loveland, T.R., Reed, B.C., Brown, J.F., Ohlen, D.O., Zhu, Z., Yang, L., Merchant, J.W., 2000. Development of a global land cover characteristics database and IGBP DISCover from 1 km AVHRR data. Int. J. Remote Sens. 21, 1303–1330.

56. Luyssaert, S., Jammet, M., Stoy, P.C., Estel, S., Pongratz, J., Ceschia, E., Churkina, G., Don, A., Erb, K., Ferlicoq, M., 2014. Land management and land-cover change have impacts of similar magnitude on surface temperature. Nat. Clim. Change 4, 389–393.

57. Manoli, G., Domec, J., Novick, K., Oishi, A.C., Noormets, A., Marani, M., Katul, G., 2016. Soil–plant–atmosphere conditions regulating convective cloud formation above southeastern US pine plantations. Glob. Change Biol. 22, 2238–2254.

58. Margolis, H.A., Ryan, M.G., 1997. A physiological basis for biosphere–atmosphere interactions in the boreal forest: an overview. Tree Physiol. 17, 491–499.

59. Matthes, J.H., Sturtevant, C., Verfaillie, J., Knox, S., Baldocchi, D., 2014. Parsing the variability in CH4 flux at a spatially heterogeneous wetland: Integrating multiple eddy covariance towers with high-resolution flux footprint analysis. J. Geophys. Res. Biogeosciences 119, 1322–1339.

60. Mesinger, F., DiMego, G., Kalnay, E., Mitchell, K., Shafran, P.C., Ebisuzaki, W., Jović, D., Woollen, J., Rogers, E., Berbery, E.H., 2006. North American regional reanalysis. Bull. Am. Meteorol. Soc. 87, 343–360.

61. Moeys, J., 2018. The soil texture wizard: R functions for plotting, classifying, transforming and exploring soil texture data. CRAN R-Proj.

62. Moore, C.E., Berardi, D.M., Blanc-Betes, E., Dracup, E.C., Egenriether, S., Gomez-Casanovas, N., Hartman, M.D., Hudiburg, T., Kantola, I., Masters, M.D., 2020. The carbon and nitrogen cycle impacts of reverting perennial bioenergy switchgrass to an annual maize crop rotation. GCB Bioenergy 12, 941–954.

63. Moore, C.E., von Haden, A.C., Burnham, M.B., Kantola, I.B., Gibson, C.D., Blakely, B.J., Dracup, E.C., Masters, M.D., Yang, W.H., DeLucia, E.H., 2021. Ecosystem-scale biogeochemical fluxes from three bioenergy crop candidates: How energy sorghum compares to maize and miscanthus. GCB Bioenergy 13, 445–458.

64. Mulla, D., McBratney, A.B., 2002. Soil spatial variability. Soil Phys. Companion 343373.

65. Novick, K., 2022. The science needed for robust, scalable, and credible nature-based climate solutions in the United States: Full Report.

66. Novick, K.A., Biederman, J., Desai, A., Litvak, M., Moore, D.J., Scott, R., Torn, M., 2018. The AmeriFlux network: A coalition of the willing. Agric. For. Meteorol. 249, 444–456.

67. Novick, K.A., Katul, G.G., 2020. The duality of reforestation impacts on surface and air temperature. J. Geophys. Res. Biogeosciences 125, e2019JG005543.

68. Novick, K.A., Metzger, S., Anderegg, W.R., Barnes, M., Cala, D.S., Guan, K., Hemes, K.S., Hollinger, D.Y., Kumar, J., Litvak, M., 2022. Informing Nature-based Climate Solutions for the United States with the best-available science. Glob. Change Biol.

69. Novick, K.A., Oishi, A.C., Ward, E.J., Siqueira, M.B., Juang, J., Stoy, P.C., 2015. On the difference in the net ecosystem exchange of CO 2 between deciduous and evergreen forests in the southeastern United States. Glob. Change Biol. 21, 827–842.

70. Oren, R., Hsieh, C.-I., Stoy, P., Albertson, J., Mccarthy, H.R., Harrell, P., Katul, G.G., 2006. Estimating the uncertainty in annual net ecosystem carbon exchange: Spatial variation in turbulent fluxes and sampling errors in eddy-covariance measurements. Glob. Change Biol. 12, 883–896.

71. Paleri, S., Desai, A.R., Metzger, S., Durden, D., Butterworth, B.J., Mauder, M., Kohnert, K., Serafimovich, A., 2022. Space-scale resolved surface fluxes across a heterogeneous, mid-latitude forested landscape. J. Geophys. Res. Atmospheres e2022JD037138.

72. Pastorello, G., Trotta, C., Canfora, E., Chu, H., Christianson, D., Cheah, Y.-W., Poindexter, C., Chen, J., Elbashandy, A., Humphrey, M., 2020. The FLUXNET2015 dataset and the ONEFlux processing pipeline for eddy covariance data. Sci. Data 7, 1–27.

73. Poe, J., Reed, D.E., Abraha, M., Chen, J., Dahlin, K.M., Desai, A.R., 2020. Geospatial coherence of surface-atmosphere fluxes in the upper Great Lakes region. Agric. For. Meteorol. 295, 108188.

74. Poggio, L., De Sousa, L.M., Batjes, N.H., Heuvelink, G., Kempen, B., Ribeiro, E., Rossiter, D., 2021. SoilGrids 2.0: producing soil information for the globe with quantified spatial uncertainty. Soil 7, 217–240.

75. Post, H., Hendricks Franssen, H.-J., Graf, A., Schmidt, M., Vereecken, H., 2015. Uncertainty analysis of eddy covariance CO 2 flux measurements for different EC tower distances using an extended two-tower approach. Biogeosciences 12, 1205–1221.

76. R Core Team, 2022. R: A Language and Environment for Statistical Computing.

77. Rebane, S., Jõgiste, K., Põldveer, E., Stanturf, J.A., Metslaid, M., 2019. Direct measurements of carbon exchange at forest disturbance sites: a review of results with the eddy covariance method. Scand. J. For. Res. 34, 585–597.

78. Rey-Sánchez, A.C., Bohrer, G., Morin, T.H., Shlomo, D., Mirfenderesgi, G., Gildor, H., Genin, A., 2017. Evaporation and CO2 fluxes in a coastal reef: an eddy covariance approach. Ecosyst. Health Sustain. 3, 1392830.

79. Rey-Sanchez, C., Arias-Ortiz, A., Kasak, K., Chu, H., Szutu, D., Verfaillie, J., Baldocchi, D., 2022. Detecting Hot Spots of Methane Flux Using Footprint-Weighted Flux Maps. J. Geophys. Res. Biogeosciences 127, e2022JG006977.

80. Richardson, A.D., Aubinet, M., Barr, A.G., Hollinger, D.Y., Ibrom, A., Lasslop, G., Reichstein, M., 2012. Uncertainty quantification, in: Eddy Covariance. Springer, pp. 173–209.

81. Richardson, A.D., Hollinger, D.Y., 2005. Statistical modeling of ecosystem respiration using eddy covariance data: maximum likelihood parameter estimation, and Monte Carlo simulation of model and parameter uncertainty, applied to three simple models. Agric. For. Meteorol. 131, 191–208.

82. Rohatyn, S., Rotenberg, E., Tatarinov, F., Carmel, Y., Yakir, D., 2022. Large variations in afforestation-related climate cooling and warming effects across short distances. bioRxiv 2022–09.

83. Runkle, B.R., Rigby, J.R., Reba, M.L., Anapalli, S.S., Bhattacharjee, J., Krauss, K.W., Liang, L., Locke, M.A., Novick, K.A., Sui, R., 2017. Delta-flux: An eddy covariance network for a climate-smart lower Mississippi Basin. Agric. Environ. Lett. 2, ael2017-01.

84. Runkle, B.R., Suvočarev, K., Reba, M.L., Reavis, C.W., Smith, S.F., Chiu, Y.-L., Fong, B., 2018. Methane emission reductions from the alternate wetting and drying of rice fields detected using the eddy covariance method. Environ. Sci. Technol. 53, 671–681.

85. Schimel, D., Hargrove, W., Hoffman, F., MacMahon, J., 2007. NEON: A hierarchically designed national ecological network. Front. Ecol. Environ. 5, 59–59.

86. Seddon, N., 2022. Harnessing the potential of nature-based solutions for mitigating and adapting to climate change. Science 376, 1410–1416.

87. Shveytser, V., Stoy, P.C., Butterworth, B.J., Wiesner, S., Skaggs, T., Murphy, B., Wutzler, T., El-Madany, T.S., Desai, A.R., 2022. Evaporation and transpiration from multiple proximal forests and wetlands. Authorea Prepr.

88. Slessarev, E., Lin, Y., Bingham, N., Johnson, J., Dai, Y., Schimel, J., Chadwick, O., 2016. Water balance creates a threshold in soil pH at the global scale. Nature 540, 567–569.

89. Starr, G., Staudhammer, C.L., Wiesner, S., Kunwor, S., Loescher, H.W., Baron, A.F., Whelan, A., Mitchell, R.J., Boring, L., 2016. Carbon dynamics of Pinus palustris ecosystems following drought. Forests 7, 98.

90. Stoy, P.C., Katul, G.G., Siqueira, M.B., Juang, J., Novick, K.A., McCarthy, H.R., Christopher Oishi, A., Uebelherr, J.M., Kim, H., Oren, R., 2006. Separating the effects of climate and vegetation on evapotranspiration along a successional chronosequence in the southeastern US. Glob. Change Biol. 12, 2115–2135.

91. Stoy, P.C., Katul, G.G., Siqueira, M.B., Juang, J.-Y., Novick, K.A., McCarthy, H.R., Oishi, A.C., Oren, R., 2008. Role of vegetation in determining carbon sequestration along ecological succession in the southeastern United States. Glob. Change Biol. 14, 1409– 1427.

92. Stoy, P.C., Khan, A.M., Wipf, A., Silverman, N., Powell, S.L., 2022. The spatial variability of NDVI within a wheat field: Information content and implications for yield and grain protein monitoring. PloS One 17, e0265243.

93. Stoy, P.C., Richardson, A.D., Baldocchi, D.D., Katul, G.G., Stanovick, J., Mahecha, M.D., Reichstein, M., Detto, M., Law, B.E., Wohlfahrt, G., 2009. Biosphere-atmosphere exchange of CO 2 in relation to climate: a cross-biome analysis across multiple time scales. Biogeosciences 6, 2297–2312.

94. Sulkava, M., Luyssaert, S., Zaehle, S., Papale, D., 2011. Assessing and improving the representativeness of monitoring networks: The European flux tower network example. J. Geophys. Res. Biogeosciences 116.

95. Sun, X., Zou, C.B., Wilcox, B., Stebler, E., 2019. Effect of vegetation on the energy balance and evapotranspiration in tallgrass prairie: A paired study using the eddy-covariance method. Bound.-Layer Meteorol. 170, 127–160.

96. Tuovinen, J.-P., Aurela, M., Hatakka, J., Räsänen, A., Virtanen, T., Mikola, J., Ivakhov, V., Kondratyev, V., Laurila, T., 2019. Interpreting eddy covariance data from heterogeneous Siberian tundra: land-cover-specific methane fluxes and spatial representativeness. Biogeosciences 16, 255–274.

97. Turner, J., Desai, A.R., Thom, J., Wickland, K.P., Olson, B., 2019. Wind sheltering impacts on land-atmosphere fluxes over fens. Front. Environ. Sci. 179.

98. Ueyama, M., Yamamori, T., Iwata, H., Harazono, Y., 2020. Cooling and moistening of the planetary boundary layer in interior Alaska due to a postfire change in surface energy exchange. J. Geophys. Res. Atmospheres 125, e2020JD032968.

99. Vanden Broucke, S., Luyssaert, S., Davin, E.L., Janssens, I., Van Lipzig, N., 2015. New insights in the capability of climate models to simulate the impact of LUC based on temperature decomposition of paired site observations. J. Geophys. Res. Atmospheres 120, 5417– 5436.

100. Vanderborght, J., Graf, A., Steenpass, C., Scharnagl, B., Prolingheuer, N., Herbst, M., Franssen, H.-J.H., Vereecken, H., 2010. Within-field variability of bare soil evaporation derived from eddy covariance measurements. Vadose Zone J. 9, 943–954.

101. Verma, M., Friedl, M.A., Richardson, A.D., Kiely, G., Cescatti, A., Law, B.E., Wohlfahrt, G., Gielen, B., Roupsard, O., Moors, E.J., 2014. Remote sensing of annual terrestrial gross primary productivity from MODIS: an assessment using the FLUXNET La Thuile data set. Biogeosciences 11, 2185–2200.

102. Vick, E.S., Stoy, P.C., Tang, A.C., Gerken, T., 2016. The surface-atmosphere exchange of carbon dioxide, water, and sensible heat across a dryland wheat-fallow rotation. Agric. Ecosyst. Environ. 232, 129–140.

103. Villarreal, S., Guevara, M., Allcaraz-Segura, D., Brunsell, N., Hayes, D., Loescher, H., Vargas, R., 2018. Ecosystem functional diversity and the representativeness of environmental networks across the conterminous United States. Agric. For. Meteorol. 262, 423–433.

104. Villarreal, S., Vargas, R., 2021. Representativeness of FLUXNET sites across Latin America. J. Geophys. Res. Biogeosciences 126, e2020JG006090.

105. Voglmeier, K., Six, J., Jocher, M., Ammann, C., 2020. Soil greenhouse gas budget of two intensively managed grazing systems. Agric. For. Meteorol. 287, 107960.

106. Volik, O., Petrone, R., Kessel, E., Green, A., Price, J., 2021. Understanding the peak growing season ecosystem water-use efficiency at four boreal fens in the Athabasca oil sands region. Hydrol. Process. 35, e14323.

107. Volk, J.M., Huntington, J., Melton, F.S., Allen, R., Anderson, M.C., Fisher, J.B., Kilic, A., Senay, G., Halverson, G., Knipper, K., 2023. Development of a benchmark Eddy flux evapotranspiration dataset for evaluation of satellite-driven evapotranspiration models over the CONUS. Agric. For. Meteorol. 331, 109307.

108. Vuichard, N., Papale, D., 2015. Filling the gaps in meteorological continuous data measured at FLUXNET sites with ERA-Interim reanalysis. Earth Syst. Sci. Data 7, 157–171.

109. Xiao, J., Chen, J., Davis, K.J., Reichstein, M., 2012. Advances in upscaling of eddy covariance measurements of carbon and water fluxes. J. Geophys. Res. Biogeosciences 117.

110. Yuan, K., Zhu, Q., Zheng, S., Zhao, L., Chen, M., Riley, W.J., Cai, X., Ma, H., Li, F., Wu, H., 2021. Deforestation reshapes land-surface energy-flux partitioning. Environ. Res. Lett. 16, 024014.

111. Zhang, Q., Barnes, M., Benson, M., Burakowski, E., Oishi, A.C., Ouimette, A., Sanders-DeMott, R., Stoy, P.C., Wenzel, M., Xiong, L., 2020. Reforestation and surface cooling in temperate zones: Mechanisms and implications. Glob. Change Biol. 26, 3384–3401.

